# A muscle-related contractile tissue specified by myocardin-related transcription factor activity in Porifera

**DOI:** 10.1101/2021.04.11.439235

**Authors:** J. Colgren, S.A. Nichols

## Abstract

Muscle-based movement is a hallmark of animal biology, but the evolutionary origins of myocytes – the cells that comprise muscle tissues – are unknown. Sponges (Porifera) provide an opportunity to reconstruct the earliest periods of myocyte evolution. Although sponges are believed to lack muscle, they are capable of coordinated whole-body contractions that purge debris from internal water canals. This behavior has been observed for decades, but their contractile tissues remain uncharacterized; it is an open question whether they have affinity to muscle or non-muscle contractile tissues in other animals. Here, we characterize the endothelial-like lining of water canals (the endopinacoderm) as a primary contractile tissue in the sponge *Ephydatia muelleri*. We find tissue-wide organization of contractile actin-bundles that contain striated-muscle myosin II and transgelin, and that contractions are regulated by the release of internal Ca^2+^ stores upstream of the myosin-light-chain-kinase (MLCK) pathway. Further, we show that the endopinacoderm is developmentally specified by myocardin-related transcription factor (MRTF) as part of an environmentally-inducible transcriptional complex that otherwise is known to function in muscle development, plasticity, and regeneration in other animals. We conclude that both muscle tissues and the endopinacoderm evolved from a common, multifunctional contractile tissue in the animal stem-lineage. Furthermore, as an actin-regulated force-sensor, MRTF-activity offers a mechanism for how water canals dynamically remodel in response to flow and can re-form normally from stem-cells in the absence of the intrinsic positional cues characteristic of embryogenesis in other animals.

## Introduction

Sponges are one of two animal lineages believed to lack neurons and myocytes – fundamental cell types that endow animals with the ability to move and respond to environmental stimuli. Nevertheless, even without the benefit of these cell types, sponges can undergo whole-body contractions in response to changes in water flow ^1–4^ (**FigS1**). Through ciliary action sponges draw water through internal water canals where they take-up bacterial prey and dissolved organic matter ^5^. If canals become clogged by debris, decreased flow activates sensory cells ^1^; in the laboratory this can be mimicked by the addition of Sumi ink – a suspension of carbon particles, or through mechanical agitation ^3^. The signal to contract is propagated through nitric oxide (NO) signaling, and modulated by glutamate, and gamma-aminobutyric acid (GABA) to stimulate contractile tissues that purge the system ^2,6^. The major gap in this model is that the contractile tissues themselves are uncharacterized. Are the mechanisms of contraction in sponge tissues similar to non-muscle tissues in other animals, such as the apical constriction of epithelia, or to myocytes?

Bilaterians have two general categories of myocytes – fast-contracting somatic myocytes, involved in voluntary or reactive body movements, and slow-contracting visceral myocytes, involved in organ movements and the maintenance of tissue tension ^7^. In both, contractions depend on interactions between actin filaments and type-II myosin, composed of two myosin heavy chains (MyHC), two regulatory light chains (RLCs), and two essential light chains (ESCs). Contraction speed reflects the kinetic properties of the MyHC that is expressed. Fast-contracting, striated-muscle myosin heavy chain (stMyHC) is found in skeletal and cardiac muscles of vertebrates, and smooth and striated invertebrate muscles ^8–10^. Slow-contracting, non-muscle myosin heavy chain (nmMyHC) is found in vertebrate smooth muscle, and some invertebrate visceral muscles, but also functions in non-muscle contractile contexts such as cell motility, cytokinesis, and apical constriction [reviewed in ^11^].

Myocytes can be further distinguished by their contractile regulatory mechanisms. In fast contracting myocytes, the tropomyosin/troponin C complex sterically hinders stMyHC from binding to actin, but is released by Ca^2+ 12^. Troponin C-regulation is unique to muscle tissues, but is restricted to bilaterians. A potentially more ancient mechanism is the MLCK pathway, in which cytoplasmic Ca^2+^ binds to calmodulin, activating MLCK to phosphorylate the RLC of myosin II ^13,14^. The MLCK pathway functions in both muscle and non-muscle contractile contexts, but the components of this pathway are conserved in all animals and, though untested, is presumed to be the primary regulatory mechanism in non-bilaterian muscle tissues ^9^

From a developmental perspective, myocytes can be defined according to the transcription factors involved in terminal differentiation. Distinct transcription factor combinations, termed core regulatory complexes (CoRCs), specify the cellular identity of cardiac, smooth, and striated muscles in vertebrates ^7^. CoRCs are well-conserved between species and are useful indicators of cell type homology between phylogenetically distant lineages ^15^. There were likely two myocyte CoRCs present in the bilaterian stem lineage, one for smooth/cardiac muscles (slow-muscle CoRC) and one for striated (fast-muscle CoRC)^7^. Central to both is the core interaction between MRTF and a MADS-box transcription factor – either serum response factor (SRF) or myocyte enhancer factor 2 (Mef2). The slow-muscle CoRC also includes transcription factors from GATA, NK homeobox, and Fox families and the fast-muscle CoRC includes the basic helix loop helix transcription factors MyoD and E12. Because MyoD is restricted to bilaterians ^9^, the slow-muscle CoRC is presumably more ancient.

Here, we identify the endopinacoderm as a primary contractile tissue in the freshwater sponge *Ephydatia muelleri*. We find that contractions depend upon the motor-activity of striated-muscle myosin II, that contractions are regulated by the myosin light-chain kinase (MLCK) pathway, and that development is governed by a possible slow-muscle-related CoRC that includes myocardin-related transcription factor (MRTF), Fox-family transcription factors, and SRF. Together, these data indicate a deep evolutionary link between sponge contractile tissues with muscle.

## RESULTS

### Contractile actin-bundles in the endopinacoderm contain striated-muscle myosin II

Contractions in *E. muelleri* are biphasic – incurrent canals contract, forcing water into excurrent canals and causing them to expand. Excurrent canals then contract as incurrent canals relax ^3^. In light of these dynamics, a candidate contractile tissue is the epithelial-lining of canals and other internal cavities, termed the endopinacoderm ^3,16^. The endopinacoderm is reported to express contraction-related genes ^17,18^ and contains linear actin-bundles that align between adjacent cells, exhibiting tissue-wide organization (**Figure 1b**) ^3^. To test whether linear actin-bundles shorten during contraction, we measured their length in relaxed versus contracted sponges and detected a 19.7% difference in mean length/cell (**Figure 1c**).

**Figure 1.**
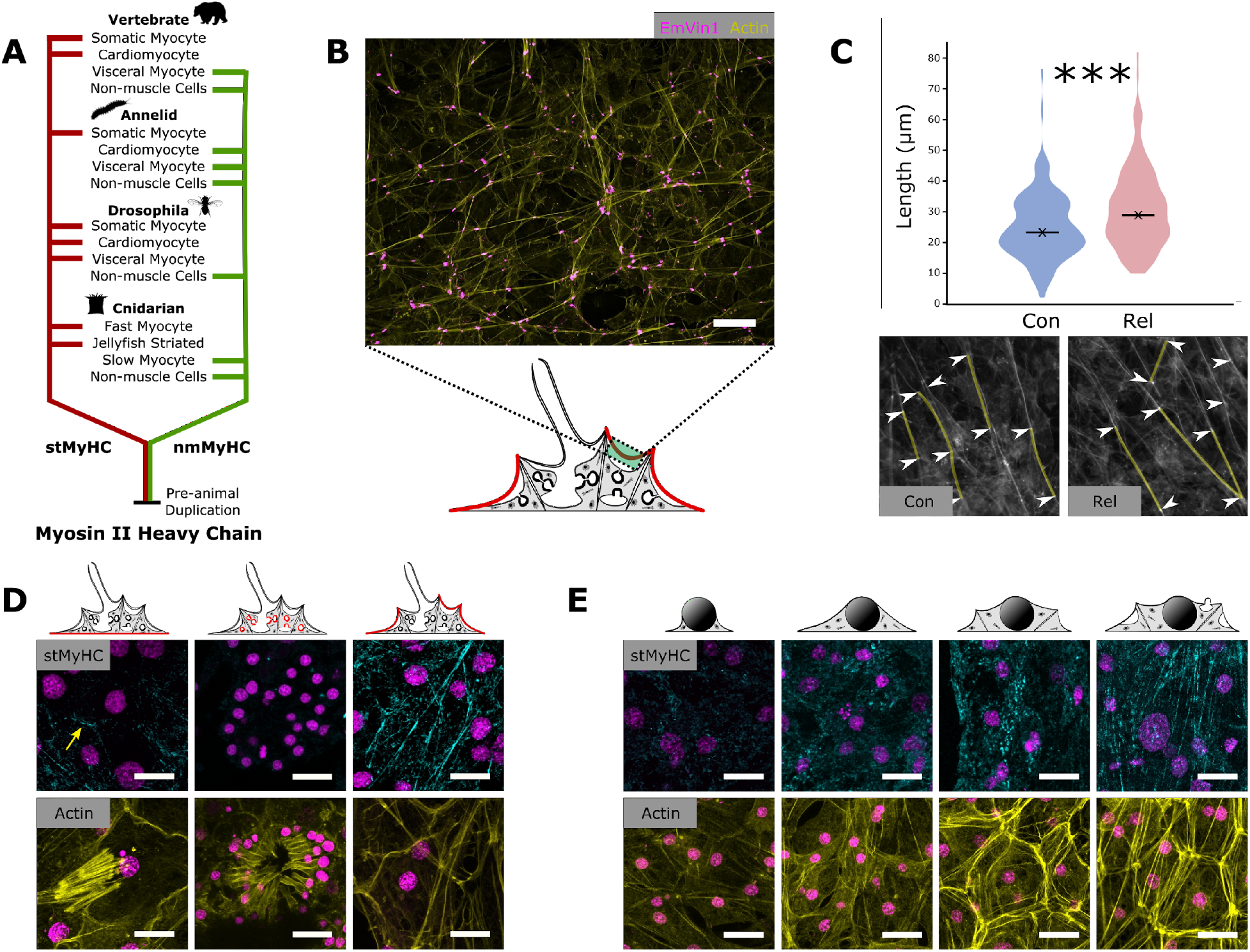
Organization of actomyosin bundles. **(A)** Myosin II Heavy Chain duplicated prior to animal origins. In modern lineages, stMyHC functions in fast-contracting myocytes, and nmMyHC functions in slow-contracting myocytes and non-muscle tissues. **(B)** Tissue-wide organization of actin-bundles (yellow) in the endopinacoderm, with cell-boundaries marked by vinculin-positive cell-cell junctions (magenta). **(C)** Actin-bundles, measured as distance between adhesion plaques, are significantly shorter in contracted tissues (p value ≪ 0.001, n=120) (top) **(D)** stMyHC immunostaining in the basopinacoderm (left), a choanocyte chamber (middle), and the apical pinacoderm (right; includes the endopinacoderm). Samples were stained for DNA (magenta), stMyHC (cyan; top), and actin (yellow; bottom). Yellow arrow shows cell boundary staining in basopinacoderm. **(E)** Developmental series of actin-bundles in the endopinacoderm stained for DNA (magenta), stMyHC (cyan; top) and actin (yellow; bottom). Scale bars 25 μm in B and 10 μm in D and E.

To test if contraction-associated shortening of actin-bundles is myosin-dependent, we immunostained tissues for stMyHC (**Figure 1d)**. Staining was diffuse during early development and increased as the endopinacoderm differentiated and canals formed. In fully-differentiated tissues, stMyHC was organized into linear structures that resembled actin-bundle orientation (**Figure 1e)**. Co-staining of actin-bundles was not compatible with the fixation conditions used for stMyHC, so to confirm their association we treated sponges with latrunculin B for increasing amounts of time prior to fixation and observed loss of stMyHC staining mirrored the loss of actin-bundles (**FigS2**).

### Contractions are regulated by the myosin light chain kinase (MLCK) pathway

Previous studies have shown that sponge contractions abate in Ca^2+^/Mg^2+^-free medium (CMFM) ^3,19^. To test whether elevated cytoplasmic Ca^2+^ can induce contractions, we treated *E. muelleri* in CMFM with the membrane-permeable calcium ionophore, ionomycin, and with the SERCA pump inhibitor, thapsigargin. Both treatments induced strong contractions (**Figure 2b**). However, because treatments were administered globally, the observed contractile response could reflect elevated Ca^2+^ in contractile tissues, or could be a secondary effect downstream of NO signalling from activated sensory cells. To decouple sensation and contraction, we treated sponges with the NO-synthase inhibitor L-NAME. As expected, L-NAME-treatment disrupted the contractile response to Sumi ink, while L-NAME treated sponges still exhibited a strong contractile response to thapsigargin (**Figure 2c**), presumably through a direct effect on the endopinacoderm.

**Figure 2.**
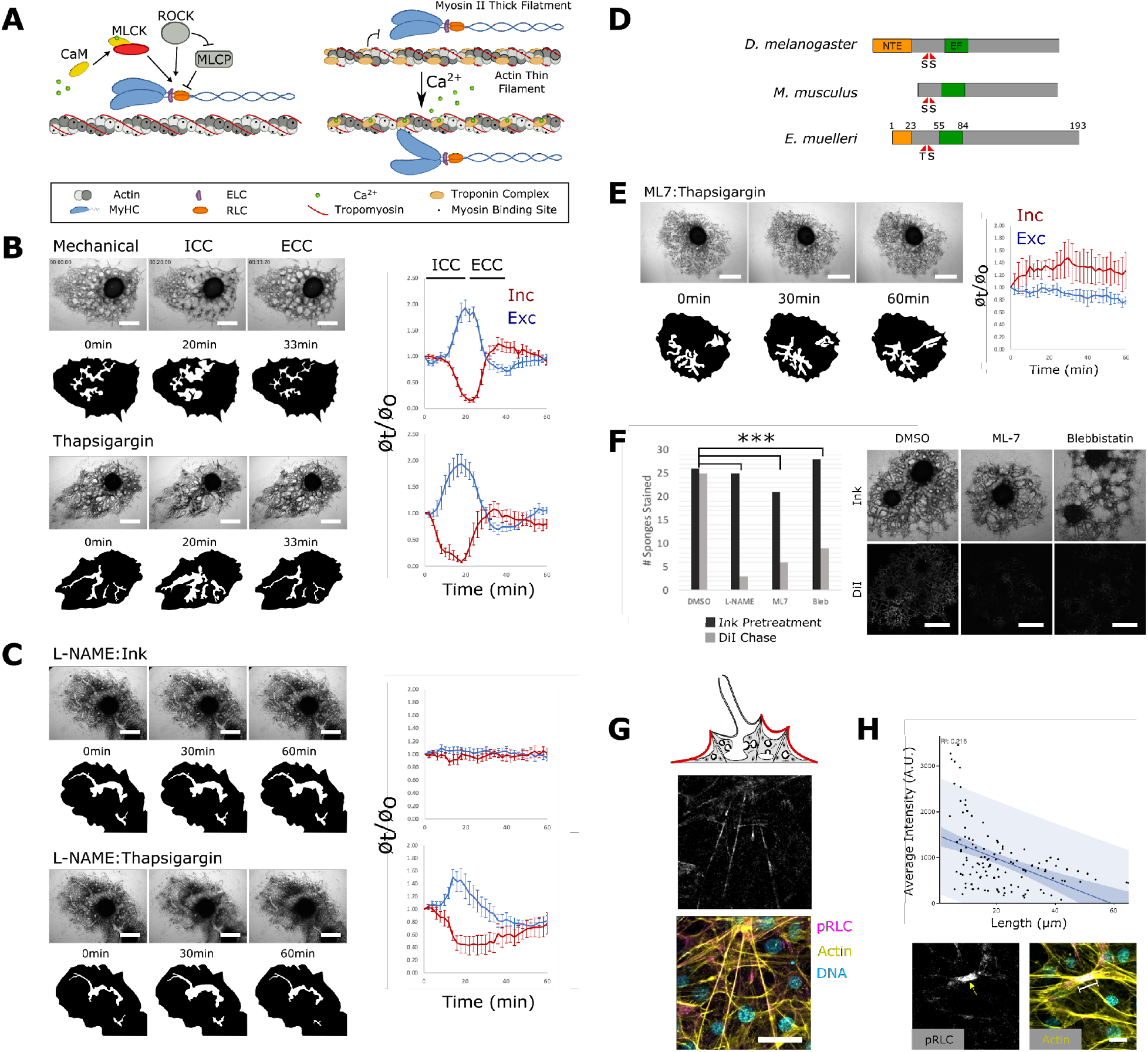
*E. muelleri* contractions are regulated by calcium-dependent MLCK activity. **(A)** In bilaterians, fast-contracting myocytes are regulated by the Troponin C/Tropomyosin complex, whereas slow-contracting myocytes are regulated by the MLCK pathway. **(B)** Time-lapse images depicting sponge contractions stimulated mechanically (top) or by treatment with 1 nM thapsigargin (bottom). Graphs (right) plot incurrent (ICC) and excurrent canal contraction (ECC) dynamics (n=6). **(C)** Top panel shows time points of a sponge treated with 20 µM L-NAME and challenged to contract with 1:500 Sumi Ink. Bottom panel shows the same sponge treated with 1 nM thapsigargin. Graphs show the average canal diameter relative to the first frame for both incurrent (red) and excurrent (blue) canals (n=6). **(D)** Predicted domain structure of the *E. muelleri* RLC homolog; a 23 amino acid N-terminal extension (NTE) exists based on location of phosphorylatable residues and EF-hand domain. **(E)** 1 nM thapsigargin had limited effect on sponges pretreated with 1 µM ML-7 to inhibit MLCK activity (n=6). **(F)** Sponges were treated with Sumi Ink to block water flow, then treated with DiI to test for restored flow as an indicator of contractile activity. The ratio of sponges that stained positive for DiI was significantly lower following treatment with 50 μg/mL L-NAME, 1 µM ML-7, and 50 μM blebbistatin compared with DMSO treated controls (Chi-square test P-value=.0007, .021, and .019 respectively). **(G)** pRLC (grayscale and magenta) immunostaining of the apical pinacoderm, counterstained with phalloidin (yellow) and Hoechst (cyan). **(H)** Scatter-plot generated by overlaying pRLC and actin images and measuring pixel intensity in the pRLC channel along the length of the actin-bundle between two adhesion plaques and simple regression analysis was performed (n=118, r^2=0.218). Below are example images with raw pRLC channel and a merged image showing a brightly pRLC-stained, short actin-bundle. Scale bars 500 μm in B, C, E, F, 10 μm in G, and 5 μm in H.

*E. muelleri* has homologs of calmodulin, MLCK, and a single RLC ortholog with conserved functional residues (**Figure 2d)**. If the release of internal Ca2+ is upstream of MLCK signaling, then contractions should be disrupted by treatment with the MLCK inhibitor ML-7. We developed a simple functional assay to test this. First, we established a concentration of Sumi ink that permanently blocked water flow in L-NAME treated sponges (i.e., sponges deficient for NO synthesis), but that could be reliably cleared by contraction of untreated, control sponges. Ink clearance and the reestablishment of flow was determined by post-treating sponges with DiI to see if it entered canals. ML-7 treated sponges were unable to efficiently clear ink and restore water flow, nor could they be induced to contract with thapsigargin **(Figure 2e)**. Similar results were obtained using blebbistatin to directly disrupt myosin ATPase activity (**Figure 2f)**.

To test for RLC phosphorylation during contraction, we immunostained sponges with a phospho-specific anti-RLC antibody (pRLC) and saw staining of contractile actin-bundles (**Figure 2g)**. Moreover, the length of individual actin-bundles was negatively correlated (R^2^=0.216, n=118) with anti-pRLC staining intensity, indicating that increased phosphorylation of the RLC is associated with actin-bundle contraction (**Figure 2h)**. Collectively, these data indicate that contraction of the endopinacoderm results from the release of internal Ca^2+^ and MLCK-phosphorylation of the RLC.

### MRTF drives endopinacoderm differentiation

MRTF, SRF, and Mef2 are broadly expressed in sponges, but expression data alone provide insufficient evidence of their possible roles in specification of contractile tissues due to MRTF’s mechanism of action. MRTF homologs are transcriptional cofactors which, when located in the cytoplasm, are inhibited through the interaction of G-actin with N-terminal RPEL (RPxxEL) repeats, ^20^; these are conserved in the *E. muelleri* ortholog (**Figure 3b**). Actin polymerization disrupts this interaction, exposing a nuclear localization signal ^21,22^. In the nucleus, MRTF can interact with SRF or Mef2 to drive the expression of contractile genes and cause myocyte differentiation ^23–26^.

**Figure 3.**
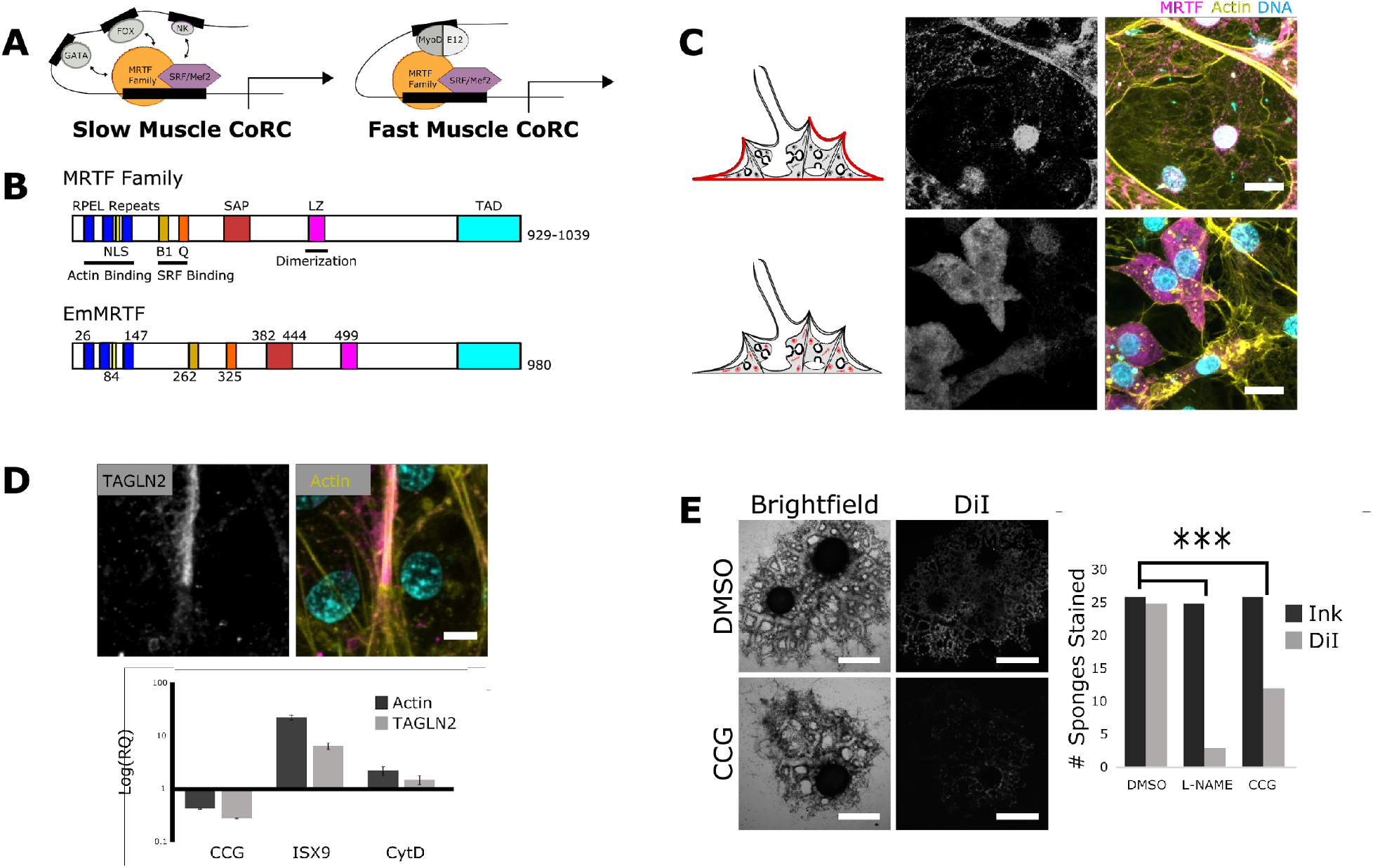
Development of contractile structures is dependent on MRTF activity. **(A)** Fast- and slow-contracting myocytes are developmentally specified by a core regulatory complex (CoRC) that includes MRTF interactions with SRF or Mef2, along with other myogenic factors unique to each CoRC. [adapted from ^7^] **(B)** Predicted domain organization of *E. muelleri* MRTF compared with vertebrate homologs (NLS; nuclear localization signal, B1; basic rich domain, SAP; SAP domain, LZ; leucine zipper, TAD; transcription activation domain). **(C)** Confocal images of pinacocytes (top) and archeocytes (bottom) immunostained for MRTF (grayscale and magenta), phalloidin (yellow), and Hoechst (cyan). **(D)** Top: confocal images of an actin-bundle immunostained for TAGNL2 (grayscale and magenta), phalloidin (yellow) and Hoechst (cyan). Bottom: qPCR showing changes to TAGNL2 levels in response to treatment with CCG-203791 (20 μM), ISX-9 (50 μM), and cytochalasin D (10 μM). **(E)** Sponges pretreated with CCG-203791 were unable to clear ink-blockages by contraction (P-value=0.004). Scale bars 500 μm E, 5 μm in C, and 2 μm in D.

We developed an antibody against the single *E. muelleri* MRTF ortholog and found that archeocytes – adult stem cells identifiable by their prominent nucleolus ^27^ – contained primarily cytosoplasmic MRTF, whereas differentiated epithelia (including the endopinacderm) contained predominantly nuclear MRTF (**Figure 3c**). Well-established targets of MRTF activity in bilaterians are members of the transgelin protein family (e.g., calponin, SM22alpha, MP20), which are often muscle specific, ^20,28^. We found that one of the three transgelin homologs in *E. muelleri*, TAGLN2, was specifically localized to contractile actin-bundles of the endpinacoderm (**Figure 3d and FigS3**). Treatment with either the MRTF inhibitor, CCG-207319, or activator, ISX-9, resulted in corollary changes in TAGLN2 expression (**Figure 3e)**. This was also seen following treatment with cytochalasin D, a potent MRTF activator through competitive binding of G-actin ^29^. Together, this suggests a conserved role for MRTF in development of the contractile-bundles of endopinacocytes as well as regulation through actin dynamics. MRTF-inhibited sponges appeared normal but had a reduced ability to contract when challenged with Sumi ink (**Figure 3f)**.

To test the effects of MRTF activation in adult stem cells, we dissociated juvenile sponges and enriched for archeocytes. We treated these cell fractions with either DMSO or ISX-9 and placed them in an attachment-free environment (this allows for the formation of primary aggregates – primmorphs – but delays differentiation). After three days, control primmorphs did not show evidence of having endopinacoderm and lacked linear actin-bundles. Primmorphs treated with ISX-9 had a more evenly spherical morphology with dense peripheral actin organized into linear bundles adjoined at adhesion plaques (**Figure 4a)**. These actin-bundles stained positive for pRLC (**FigS4**) and treated primmorphs contained elevated levels of stMyHC (**Figure 4b**). Thapsigargin had no effect on DMSO-treated aggregates but induced normal contractions in ISX-9 treated primmorphs, which exhibited a 15.0 (+/-5.5)% reduction in cross-sectional area, followed by return to resting size (**Figure 4c**).

**Figure 4.**
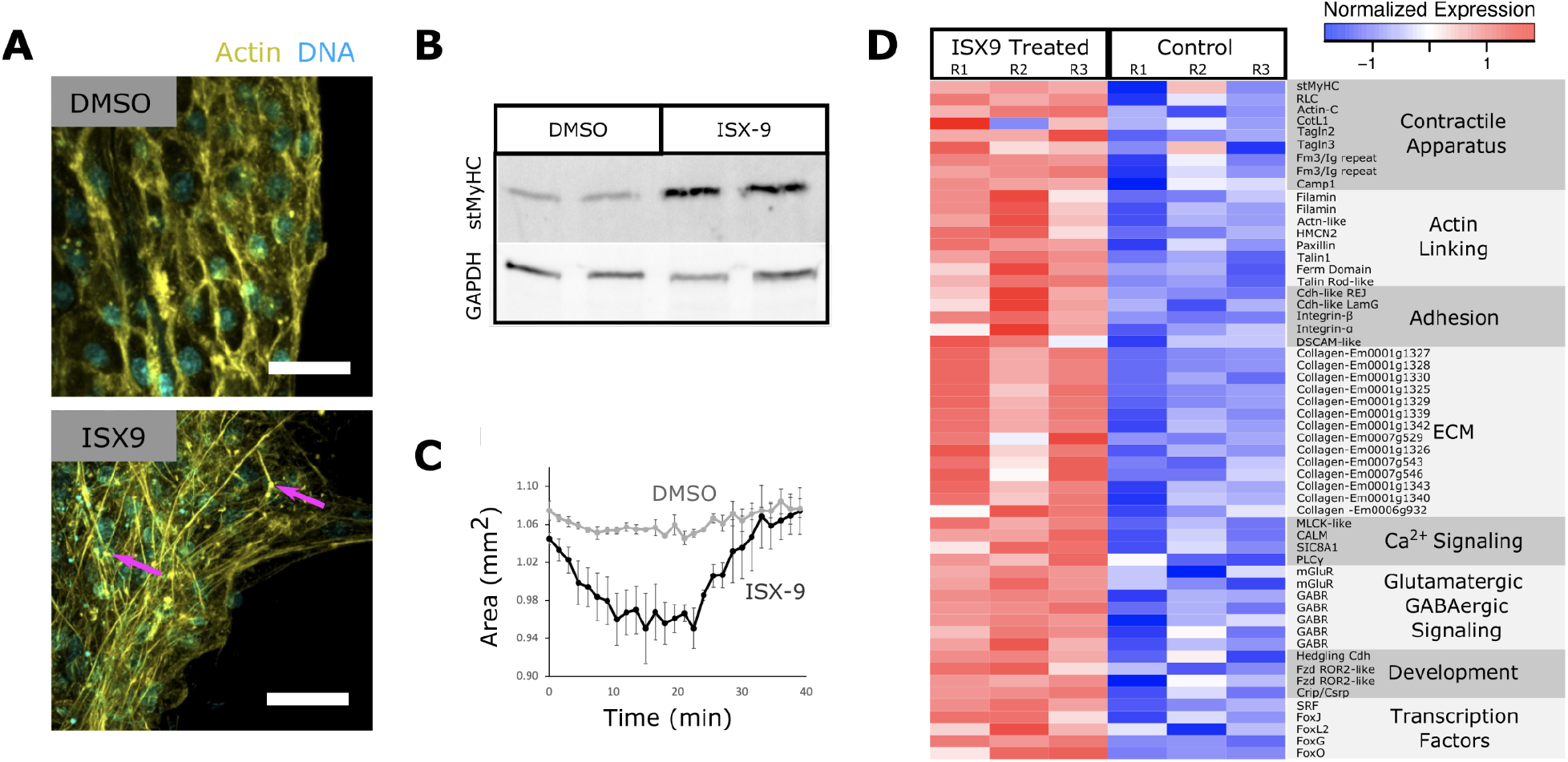
MRTF drives endopinacoderm differentiation. **(A)** Archeocyte-enriched aggregates treated with 50 µM ISX-9 or DMSO and stained with phalloidin (yellow) and Hoechst (cyan). **(B)** Western Blot showing elevated stMyHC levels in ISX-9-treated samples. **(C)** Contraction dynamics of ISX-9-treated aggregates in response to thapsigargin. **(D)** Heatmap of select transcripts that were differentially expressed in ISX-9 treated samples. Scale bars 10 μm.

### MRTF activates myogenic factors, signaling, contractile, and adhesion genes

To better understand the transcriptional response of primmorphs to MRTF activation, we sequenced mRNA from ISX-9 or DMSO treated primmorphs. Differential expression analysis of 16,712 mapped transcripts revealed that 1,390 were upregulated and 1,091 were downregulated (**FigS5**). The ISX-9 treated samples had significantly elevated expression of genes involved in contraction, cell signaling, development, and adhesion (**Figure 4d**). Contraction-related genes included stMyHC (LogFC=0.65, P=0.038), which was validated by western blot (**Figure 4b**), TAGLN2 (LogFC=1.03, P=1.0E^-3^), and components of the Ca^2+^-dependent MLCK pathway including MLCK-like serine/threonine kinase (LogFC=1.58, P=6.29E-5), calmodulin (LogFC=1.55, P=0.002), sodium/calcium exchanger 1 (SLC8A1) (LogFC=1.05 P=0.029), and Phospholipase C gamma (PLCγ) (LogFC=1.01, P=0.005) (**Figure 4d**). Upregulated signaling genes included metabotropic glutamate receptors (mGluRs) and GABA receptor subunits (GABR) (**Figure 4d**), consistent with previous studies showing a role for glutamatergic and GABAergic signaling in sponge contractile behavior ^2,6^. Upregulated developmental factors included the myogenic transcription factor SRF (LogFC=1.01, P=0.005). Four Forkhead transcription factors showed increased expression, including Fox-L2 (LogFC=1.07, P=0.008), FoxG (LogFC=1.31, P=4.16E^-5^), which is expressed in myocytes of invertebrates ^30^, FoxO, and FoxJ1 (LogFC=0.60, P=0.016 and LogFC=1.86, P=0.001 respectively). The phylogenetically broad muscle marker Crip/Csrp ^31^ was also upregulated (LogFC=1.10, P=0.001). Collectively these data support that MRTF may function as part of a slow-muscle CoRC together with SRF and Fox transcription factors.

Some of the most highly upregulated genes belonged to the collagen family, suggesting a role for the endopinacoderm in the secretion of the extracellular matrix **(Figure 4d**). As many as fourteen collagen homologs had minimal expression in control primmorphs, but high expression in ISX-9 treated samples. Adhesion molecules, including cadherins, integrins, and down syndrome cell adhesion molecule (DSCAM) had increased expression levels as well (**Figure 4d**). Though many of the upregulated genes correspond to the transcriptional profile of endopinacocytes based scRNA-seq data ^18^, upregulation of silicatein (gene cluster on scaffold 3; 1436, 1437, 1438, & 1439; LogFCs=4.42, 1.92, 4.58, & 2.20, Ps=3.22E-5, 0.018, 0.001, & 0.01 respectively), a sclerocyte marker ^18^, suggests that ISX-9 treatment caused differentiation of other cell types as well, directly or indirectly.

## DISCUSSION

Here, we establish that the endopinacoderm is a primary contractile tissue in *E. muelleri*, its contractions depend upon MLCK regulation of striated-muscle myosin II, and it is developmentally specified by MRTF-activity as part of a possible slow-muscle-related CoRC. By each of these metrics, the endopinacoderm falls within the range of variation exhibited by myocytes in other animals. For example, the ascidian *Ciona robusta* and the annelid *Platynereis dumerilii* both have striated myocytes that express stMyHC, which are regulated by the troponin C/tropomyosin complex, and are developmentally specified by a fast-muscle CoRC as well as smooth muscles that express nmMyHC and are patterned by a slow-muscle CoRC ^7,32^. The flatworm *Schistosoma mansoni*, has smooth muscles that express stMyHC, are regulated by MLCK phosphorylation of the RLC, and patterned by a fast-muscle CoRC that includes MyoD ^33,34^. Cnidarians have both fast- and slow-contracting myocytes that express either stMyHC or nmMyHC, but they lack other features of bilaterian fast-contracting muscles such as troponin C/tropomyosin regulation and the fast-muscle CoRC components MyoD ^9^. Ctenophore myocytes predominantly have smooth ultrastructure, exclusively express stMyHC, and lack troponin C/tropomyosin regulation and MyoD ^9,35^.

Our interpretation of the endopinacoderm as being muscle-related is consistent with single-cell sequencing data from the related freshwater species, *Spongilla lacustris*, which indicate that pinacocytes (data for the endopinacoderm were not disaggregated from other pinacodermal tissues) and another cell type, myopeptidocytes, cluster with myocytes from other animals ^18,30^. Myopeptidocytes are solitary cells in the interstitial extracellular matrix (mesohyl) and express contractile genes including nmMyHC, but nothing else is yet known about their developmental specification, contractile behavior, or possible regulatory mechanisms. Single-cell RNA sequencing of the demosponge *Amphimedon queenslandica* also revealed co-expression of key components of the actin-based contractile apparatus (including stMyHC) in pinacocytes ^17^.

However, to say that the endopinacoderm is muscle-related is not to assert outright, one-to-one homology of these tissues. Invertebrate muscles are often multifunctional. The epitheliomuscles of *Hydra* function in contraction, the formation of an epithelial barrier, innate immunity, and regeneration ^36^. In the planarian *Schmidtea mediterranea*, muscle also acts as a connective tissue that secretes ECM proteins (including 19 collagens) and signaling molecules that provide positional cues for regeneration from neoblasts ^37^. Similarly, the sponge endopinacoderm forms an endothelial-like barrier to the environment, is capable of phagocytosis ^38^, and expresses genes involved in sensation, metabolism, and defense ^18^. In an intriguing parallel with *S. mediterranea*, we found that MRTF-activation of endopinacoderm differentiation was associated with upregulation of ECM, including 14 collagen genes. Collectively, these data support that muscle tissues were ancestrally multifunctional and that narrowly specialized muscles with dedicated contractile functions may have evolved later.

The discovery of a muscle-related contractile tissue in *E. muelleri* has potentially important implications with respect to the timing of myocyte evolution. Depending on the phylogenetic position of sponges, which is contentious ^39^, the long-held view that they lack muscles has been interpreted as evidence that myocytes evolved after sponges diverged from other animals, or that myocytes were secondarily lost in sponges ^40^. In light of our results, the most parsimonious interpretation is that muscle-related contractile tissues predate modern animals, irrespective of the phylogenetic placement of sponges. This interpretation fits well with studies showing that the stMyHC and nmMyHC split occurred in the holozoan stem lineage ^9,41^, and at least one species of choanoflagellate – the sister group to animals – forms multicellular colonies that resemble a polarized epithelium with coordinated contractile behaviors ^42^.

Finally, the role of MRTF in endopinacoderm differentiation helps to explain the plasticity and regenerative capacity of sponge tissues. Evidently without the need for the intrinsic positional cues characteristic of embryogenesis, sponges can develop normally from archeocyte-enriched aggregates and from gemmules, and adult tissues can remodel in response to flow dynamics ^43,44^, ^45,46^. In vertebrates, MRTF is an actin-regulated force-sensor involved in muscle plasticity and regeneration ^21,47,48^, and our data indicate that similar mechanisms are operating in *E. muelleri*. We found that MRTF is located in the cytoplasm in archeocytes but is nuclear in differentiated tissues. Also, substrate attachment (or pharmacological MRTF activation) is required for endopinacoderm differentiation in archeocyte-enriched aggregates. These data support a model in which MRTF-mediated environmental feedback mechanisms drive the development of the endopinacoderm. We speculate that this could be the rule rather than the exception in sponges. Even during sexual reproduction, there is limited correlation between embryonic patterning and adult tissue identity ^49^. It is conceptually plausible that this reflects the ancestral condition and that the evolution of animal developmental mechanisms may have involved a transition in which ancient environmental feedback mechanisms were harnessed by genetically-encoded, intrinsic patterning mechanisms ^50^.

## Conclusions

The answer to the question of whether sponge contractile tissues can be interpreted as muscle-related is nuanced. Certainly, the endopinacoderm has more in common with myocytes than with non-muscle contractile tissues in that it expresses stMyHC, has internal Ca^2+^ stores, and exhibits tissue-wide actomyosin organization. Moreover, MRTF-activation of endopinacoderm differentiation leads to the upregulation of SRF and Fox-family transcription factors, indicating conserved elements of an ancient slow-muscle CoRC. Like many invertebrate muscle tissues, the endopinacoderm is multifunctional, but unlike muscle tissues in other animals, its development may be predominantly regulated by environmental feedback mechanisms rather than intrinsic patterning.

## Acknowledgments

The authors would like to thank Jennyfer Mora Mitchell for her help with experimental design and field collection of *E. muelleri*, and Pawel Burkhardt for providing feedback on the initial preprint version of the manuscript.

## Funding

JC and SAN were supported by research grants to SAN from the National Aeronautics and Space Administration (16-EXO16_2-0041) and the National Science Foundation (IOS:2015608).

## Author contributions

JC: conceptualization, investigation, visualization, analysis, and writing. SAN: conceptualization, supervision, project administration, and writing.

## Competing interests

The authors declare no competing interests.

## Data and materials availability

Raw RNAseq reads have been deposited at the NCBI SRA (accession number/BioProject PRJNA718521, BioSamples SAMN18537458, SAMN18537459, SAMN18537460, SAMN18537461, SAMN18537462, SAMN18537463).

## Supplementary Materials

**File 1** - Supplemental Figures S1-S12

**File 2** - Protein sequences used in study

**Video 1** - Time lapse *E. muelleri* mechanical contraction

**Video 2** - Time lapse *E. muelleri* thapsigargin contraction

**Video 3** - Time lapse *E. muelleri* L-NAME treated with ink

**Video 4** - Time lapse *E. muelleri* L-NAME treated with thapsigargin

**Video 5** - Time lapse *E. muelleri* ML-7 treated with thapsigargin

## Notes

### Competing Interest Statement

The authors have declared no competing interest.

### Summary of Updates

Minor formatting, text, and figure alterations.

