## Supplementary File 1 - Figures for "A muscle-related contractile tissue specified by myocardin-related transcription factor activity in Porifera"

**A**

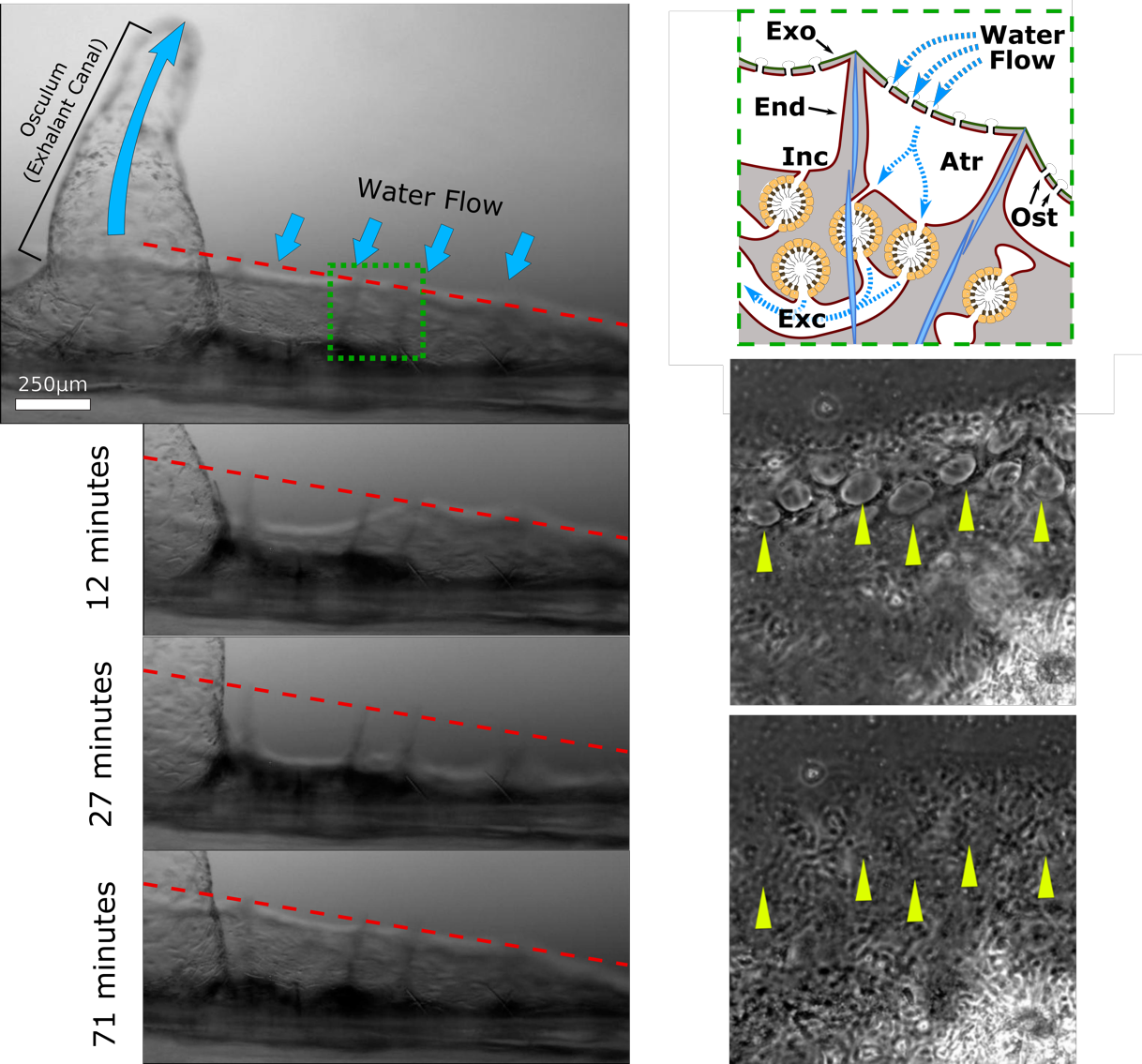

**B**

**C**

**S1. Sponges undergo coordinated whole-body contractions.** **(A)** Time-lapse series of a contraction in Ephydatia muelleri. The anatomy of the region shown to contract is illustrated in panel **(B).**The tent-like outer layer is supported by spicules (light blue) and is composed of two epithelia, exopinacoderm (Exp) and endopinacoderm (Enp) which house a thin extracellular matrix containing migratory cells. Water is drawn into the atrium (Atr) through incurrent pores (ostia; Ost) and into choanocyte chambers (orange) then into excurrent canals (Exc) towards the osculum, where it exits the sponge. The ostia on the surface (top) close upon contraction (bottom), which allows for internal water pressure to increase, clearing debris blockages.

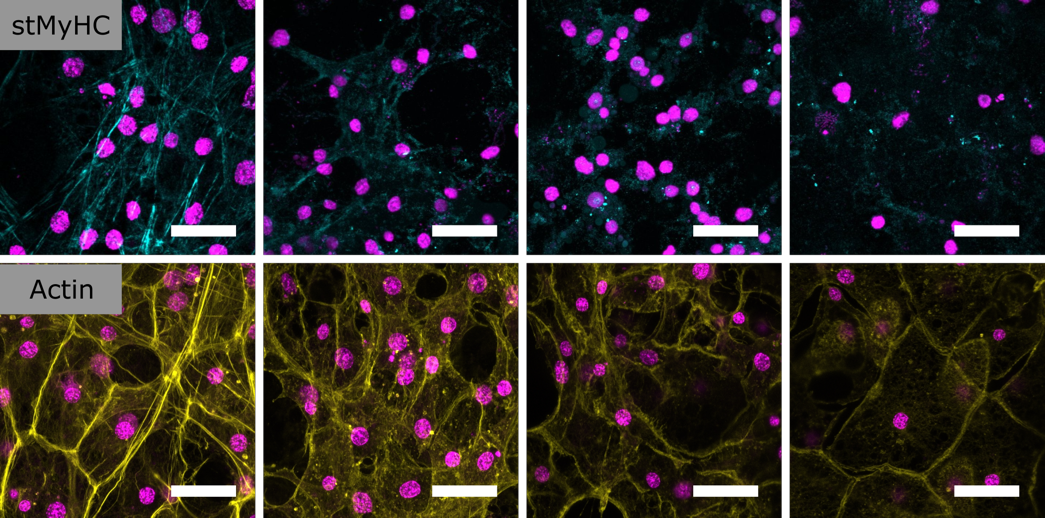

0 mins

5 mins

10 mins

20 mins

**S2. Loss of stMyHC signal mirrors actin bundles following treatment with Latrunculin B.** Juvenile sponges were treated with 20μM Latrunculin B for increasing amounts of time prior to fixation. stMyHC signal (cyan; top panel) can be seen to become more diffuse and less organized between 5 and 10 minutes following treatment. This is consistent with the loss of the actin bundles of the endopinacoderm visualized by phalloidin staining (yellow; bottom panel). DNA shown in magenta. Scale bars 20μm.

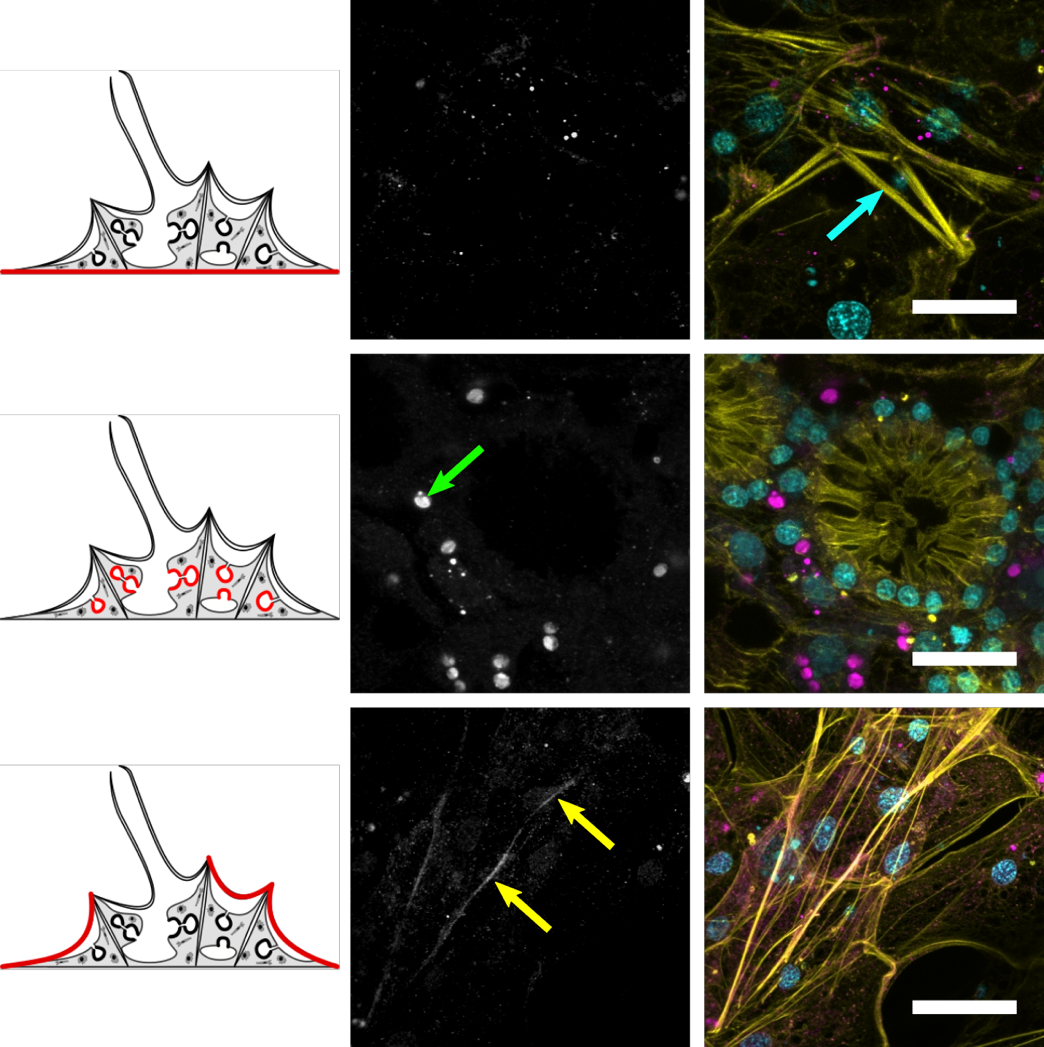

**S3. Staining for EmTAGLN2 is restricted to actin bundles of the endopincoderm.** Maximum intensity projections of confocal stacks through the basopinacoderm (top), choanoderm (middle), and apical pinacoderm (bottom). Raw antibody channel shown in gray next to merge images showing DNA (cyan), actin (yellow), and EmTAGLN2 (magenta). Cyan arrow shows large stress fiber-like structure in a basopinacocytes, which does not stain for EmTAGLN2. Green arrow highlights auto-fluorescent aglae symbiont, and yellow arrow highlights EmTAGLN2 staining along linear filaments. Scale bars, 10μm.

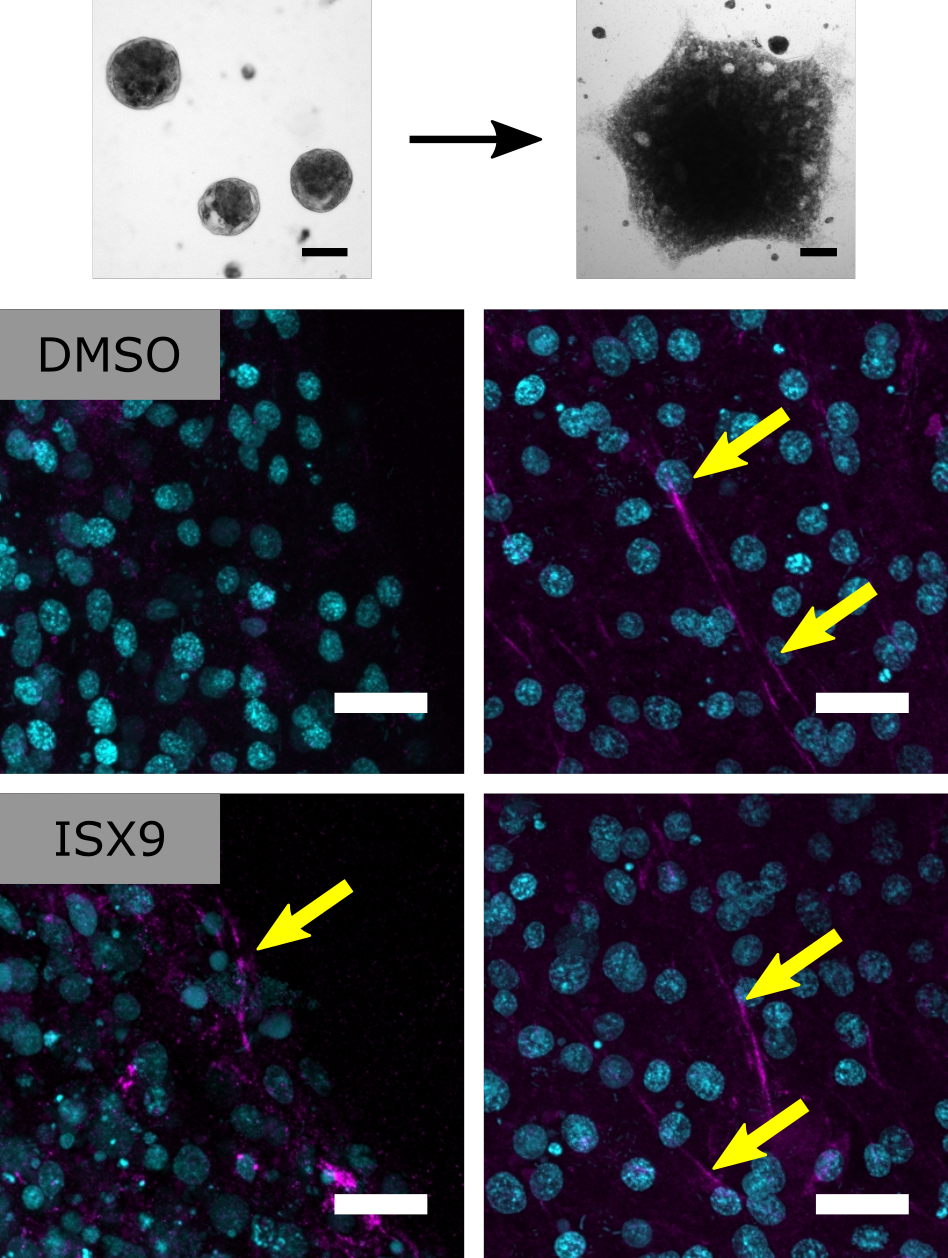

**S4. Primmorphs treated with ISX9 stain for pRLC, suggesting a contractile phenotype.** Top panel shows brightfield images of primmorphs (left) and a newly attachted sponge (right). Middle panel shows pRLC staining (magenta) in a control, day 3 primmorph (left) and a new attached sponge (right). Signal along actin bundles (yellow arrows) becomes visible following attachment. Bottom panels shows a day 3 primmorph treated with ISX9 (left) and a treated sponge following attachment (right). pRLC signal is visible along linear structures in the treated primmorphs. DNA in cyan. Scale bars 100μm and 10μm.

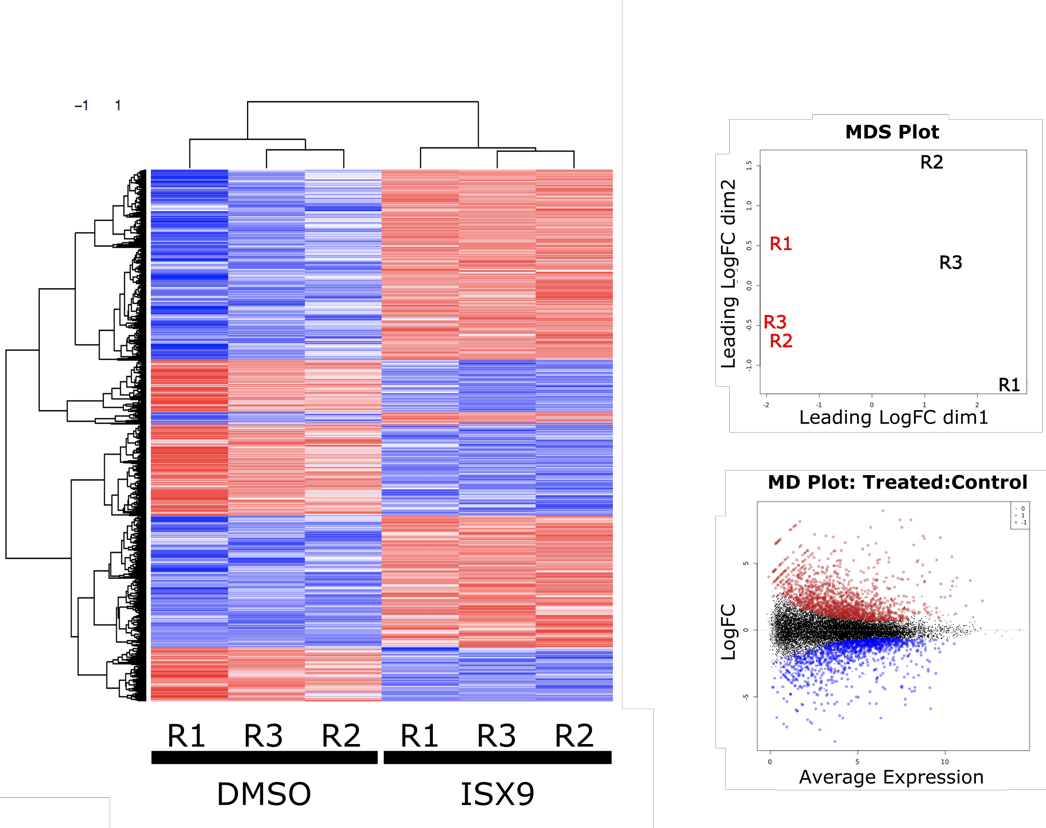

**A**

**B**

**C**

**S5. Treatment shows a large and consistent transcriptional response in primmorphs. (A)** Heatmap of full list of differentially expressed transcripts between treated and control sponges. Red shows upregulation in treated relative to control, while blue show decreased expression. Rows and columns grouped based on clustering of normalized expression. **(B)** MSD plot for replicates of ISX9 treated (red) and DMSO (black) primmorphs. **(C)** MD plot for differentially expressed transcripts between treatment and control. Red shows significant increased expression in treatment and blue show significant decreased expression.

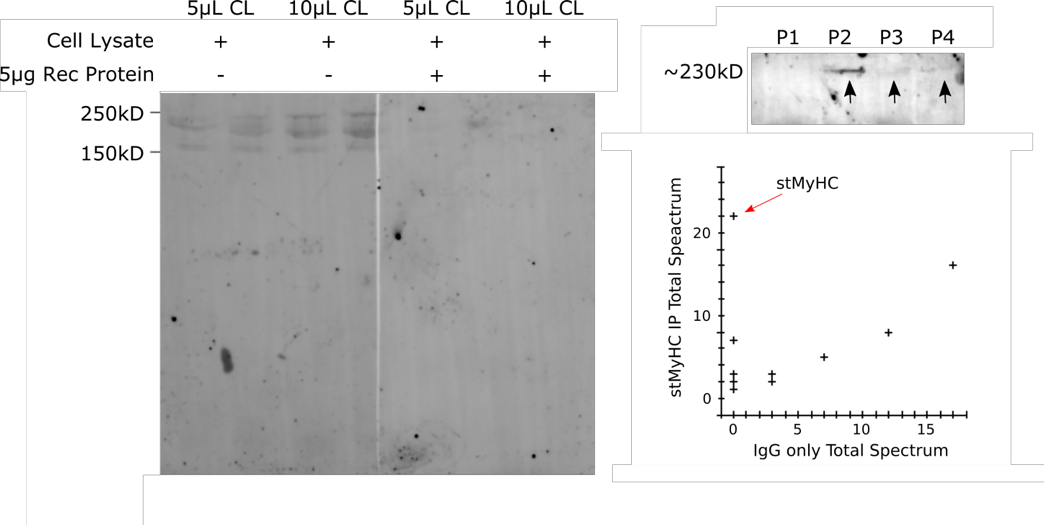

**A**

**B**

**C**

**S6. Validation of custom EmstMyHC antibody. (A)** Western blot of sponge whole cell lysates shows band at correct size for the predicted protein, which is lost following competition with recombinant peptide antigen. **(B)** Western blot of immunoprecipitation precipitates shows protein present in elutions 2-4. **(C)** Results for mass spectrometry performed on stMyHC precipitates and IgG control precipitates show stMyHC as strongest unique hit.

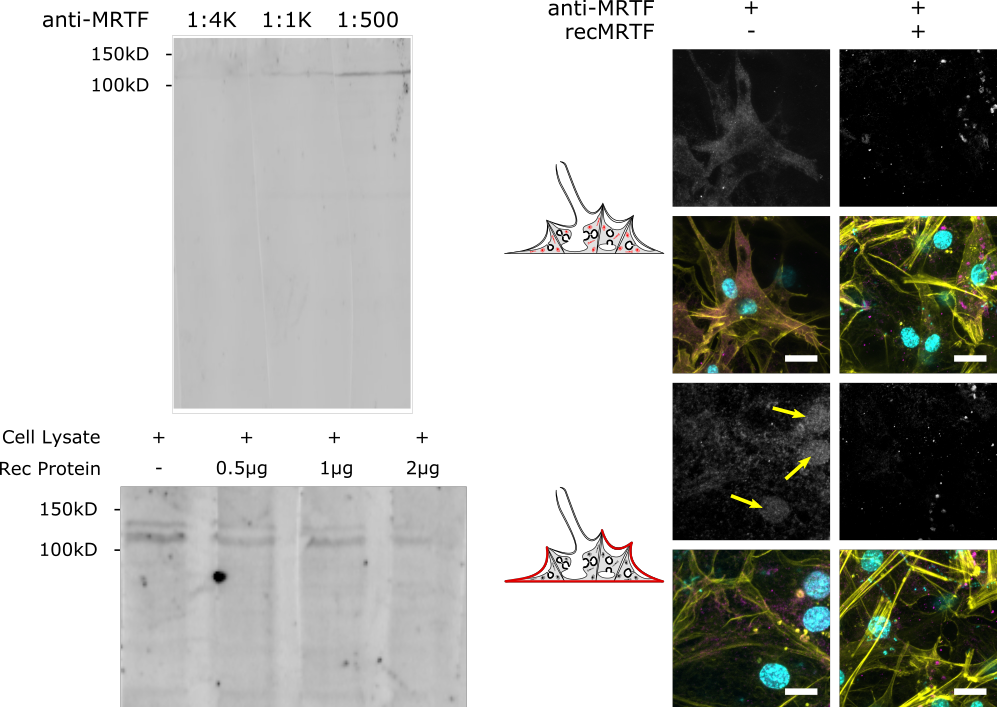

**A**

**B**

**C**

**S7. Validation of custom EmMRTF antibody. (A)** Western blot of sponge whole cell lysates using increasing concentrations of antibody shows a specific band at the predicted MW of the protein. **(B)** Western blot of sponge whole cell lysates blotted with anti-EmMRTF competed with increasing concentration of recombinant protein shows a loss in signal. **(C)** Immunofluorescent images of juvenile sponges stained with anti-EmMRTF either without recombinant protein (left) or with (right). Top portion shows migratory cells in the mesohyl and bottom portion shows pinacocytes. Grayscale images of raw antibody channel (top) and merged imagines show actin (yellow), EmMRTF (magenta), and DNA (cyan). Both broad cytoplasmic staining of migratory cells and nuclear staining (yellow arrows) of pinacocytes is lost following competition with recombinant protein. Scale bars 5μm.

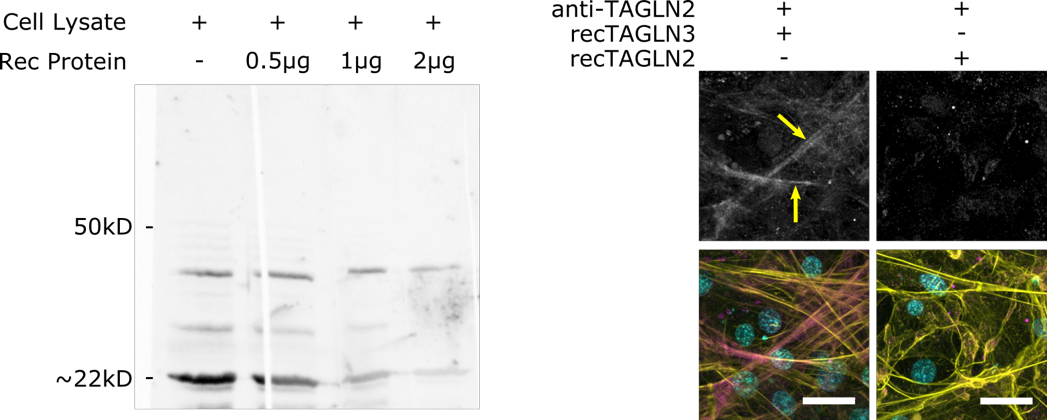

**A**

**B**

**S8. Validation of custom EmTAGLN2 antibody. (A)** Western blot of sponge whole cell lysates

shows strong band at the predicted size of the protein, which is lost with increasing concentration of recombinant protein. Higher MW bands are consistent with patterns seen for western blots of transgelins in other animals. **(B)** Immunofluorescent images of sponges stained with anti-EmTAGLN2 (magenta) competed with recombinant EmTAGLN3 (left) or recombinant EmTAGLN2 (right). Counter stained for actin (yellow) and DNA (cyan). Raw antibody channel (top) shows signal remains at actin bundles following competition with EmTAGLN3 (yellow arrows) but is lost following competition with EmTAGLN2. Scale bars 10 μm.

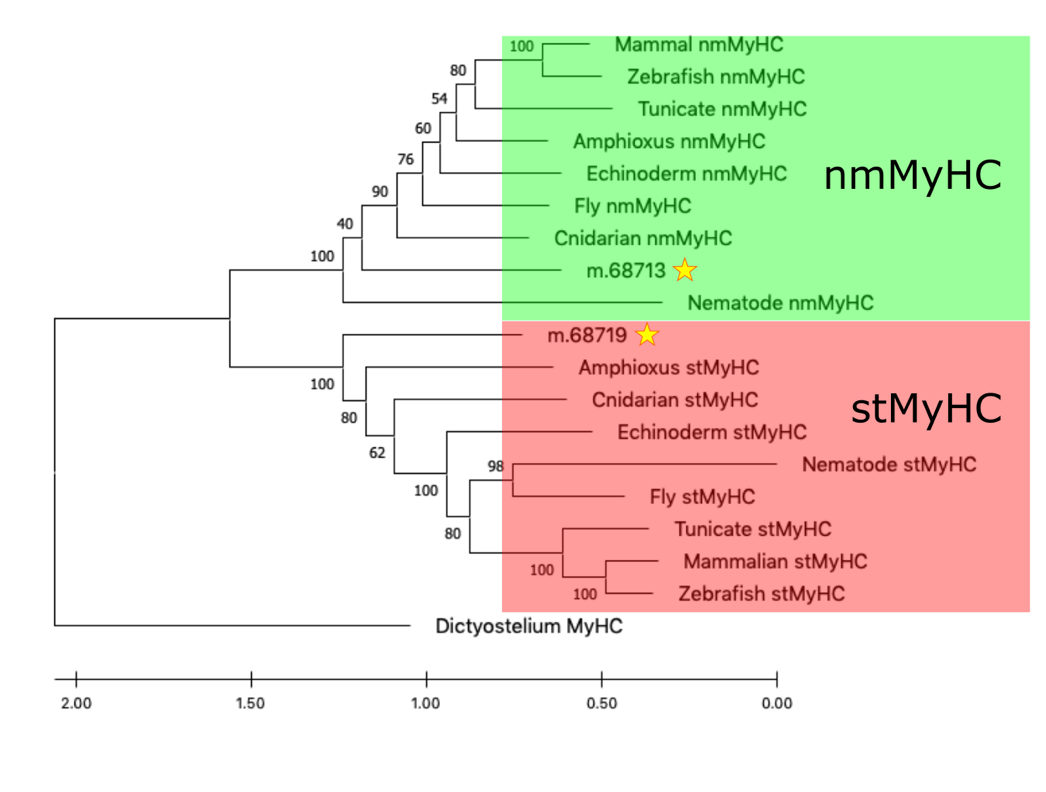

**S9. *E. muelleri* has a nmMyHC and stMyHC.** The type II MyHC proteins found in *E. muelleri* (labeled with stars) fall into either stMyHC clade or nmMyHC clade. Phylogeny determined by maximum likelihood method. Tree rooted with MyHC from *Dictyostelium.* Support values represent 1000 bootstrap iterations.

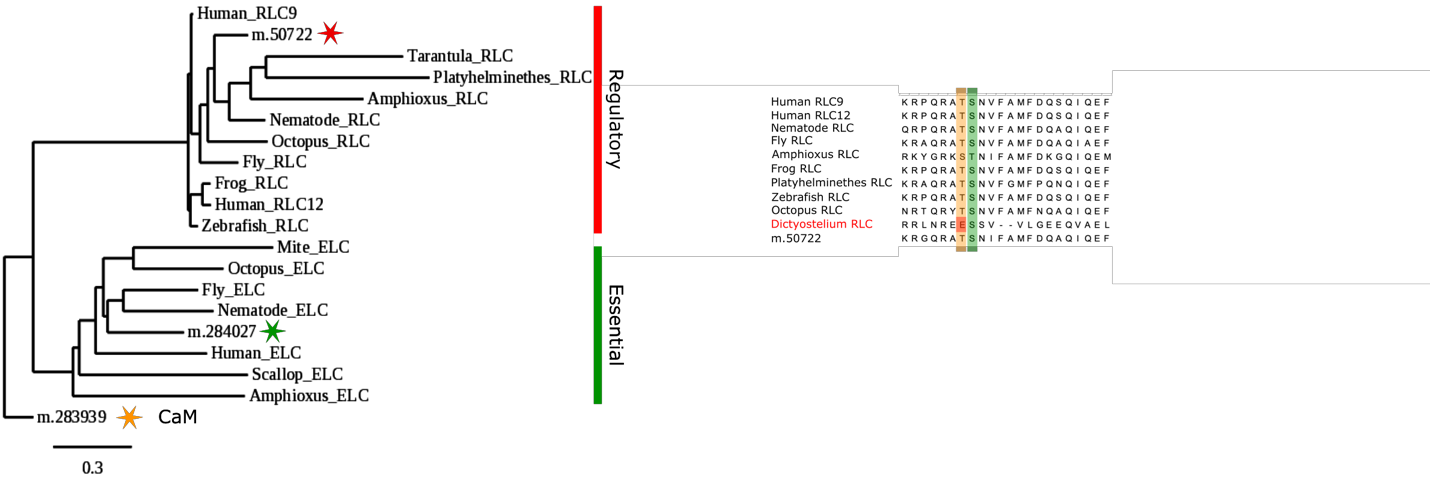

**B**

**A**

**S10. *E. muelleri* orthologs of RLC and ELC.** **(A)** Maximum likelihood phylogeny of evolutionarily related RLC and ELC rooted with the sponge EF-hand protein calmodulin (CaM). *E. muelleri* has an ortholog in the regulatory light chain group (red star) and the essential light chain group (green star). **(B)** Alignment of the region around the MLCK phosphorylation site showing conservation of phosphorylatable residues (highlighted in orange and green) and flanking regions in all animals including *E. muelleri* (m.50722).

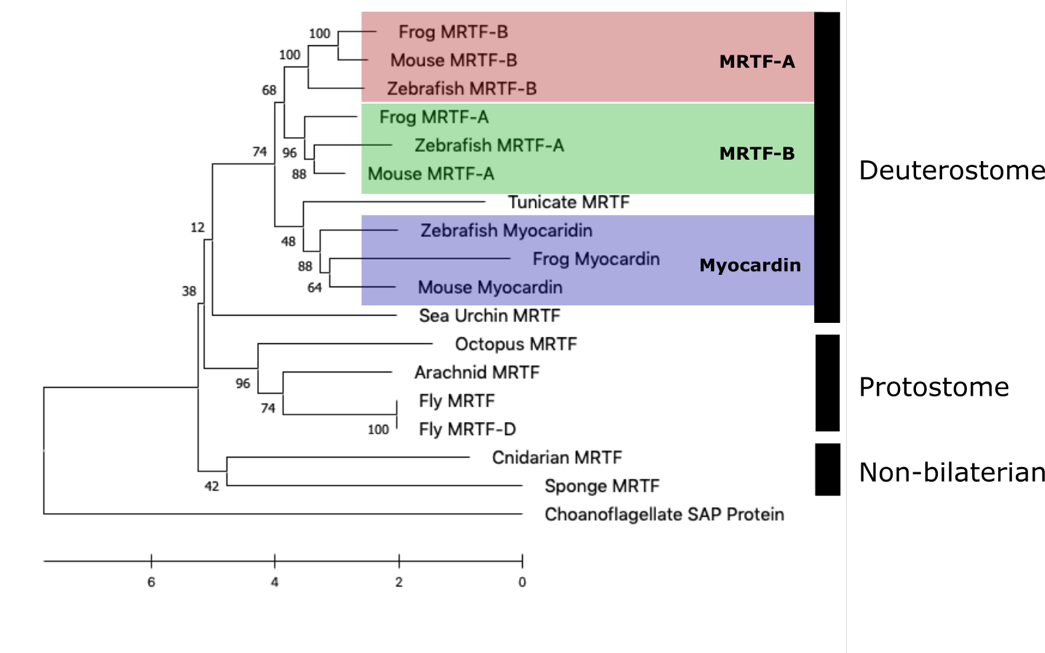

**S11. MRTF family underwent expansion in vertebrate lineage.** Maximum likelihood phylogeny of MRTF family proteins from a variety of animals, which closely matches their phylogenetic positioning. Single orthologs are generally present for non-bilaterians, protostomes, echinoderms, and hemichordates while vertebrate proteins can be divided into MRTF-A, MRTF-B, and myocardin. Choanoflagellate SAP domain containing protein was used as an outgroup. Support values based on 100 bootstrap iterations.

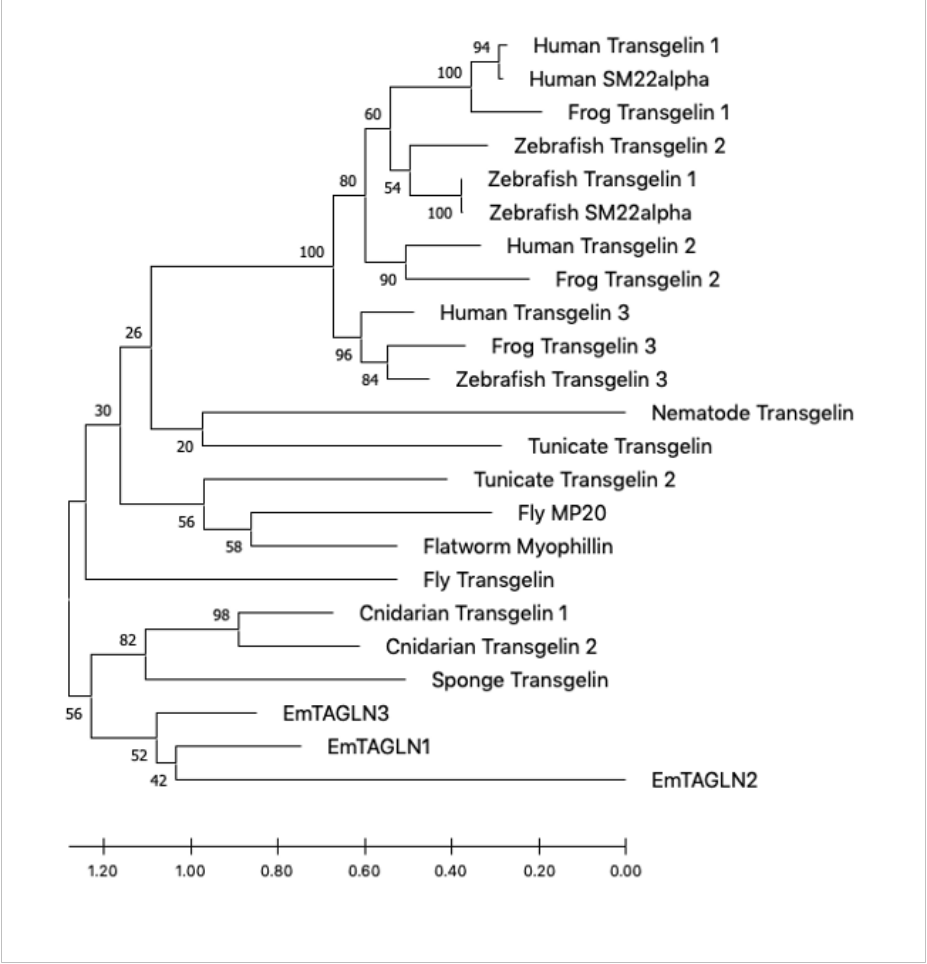

**S12. *E. muelleri* transgelins represent a lineage specific expansion**. Maximum likelihood phylogeny of transgelin family proteins from a variety of animals. Proteins grouping fit nicely with the phylogenetic positions of the animal source, which suggests an expansion prior to the vertebrate radiation as well as in non- bilaterian lineages. Support values are based on 100 bootstrap iterations.
